# Taxol acts differently on different tubulin isotypes

**DOI:** 10.1101/2023.02.07.527540

**Authors:** Yean Ming Chew, Robert A. Cross

## Abstract

Taxol is a critically important cancer drug that stabilises microtubules. We report that taxol acts differently on different metazoan tubulin isotypes. 50 nM taxol blocks catastrophe of human or zebrafish α1β4 but has no effect on human α1β3 microtubules. 500 nM taxol blocks catastrophe in both α1β3 and α1β4 microtubules but introduces kinks only into α1β4 microtubules. Taxol washout relaxes the kinks, suggesting taxol expands α1β4 but not α1β3 lattices. Kinesin-driven microtubule gliding detects this conformational shift - α1β4 microtubules glide at ~450 nm/sec in 400 nM taxol, but at ~750 nm/sec in 10 μM taxol, whereas α1β3 microtubules glide at ~450 nm/sec, even in 10 μM taxol. Thus, taxol readily stabilises α1β4 GDP-tubulin lattices and shifts them to a fast-gliding conformation, but stabilises α1β3 lattices much less readily and without shifting their conformation. These isotype-specific actions of taxol may drive the switch to β3 tubulin commonly seen in taxol-resistant tumours.

---

Microtubules are fundamental to eukaryotic life. They are essential for a wide variety of cellular functions, including chromosome segregation, directed vesicle and organelle transport, cell motility and the establishment of cell polarity. Microtubules self-assemble from tubulin heterodimers. In the presence of GTP, αβ tubulin heterodimers polymerise head-to-tail to form protofilaments and side-by-side to form a closed tube. Tubulation produces a stiff structure, allowing microtubules to define and support cell shape. The head-to-tail polymerisation of tubulin molecules gives microtubules an intrinsic polarity, whereby each microtubule has a fast growing plus end that displays β tubulin and a slow growing minus end that displays α tubulin (**Fig.1a**). At the ends of growing microtubules, GTP-tubulin heterodimers are captured from solution and incorporated into the tip lattice. The GTP-tubulin tip lattice is structurally stable but converts continuously via GTP hydrolysis and Pi release into the GDP-tubulin core lattice, which is structurally unstable. Competition between GTP-tubulin recruitment and its coupled conversion to GDP-tubulin leads to dynamic instability^1^, whereby the steady growth of each microtubule is interrupted by catastrophe events. Catastrophes are caused by stochastic breaches in the GTP-tubulin cap (**Fig. 1a**, left) that expose the underlying GDP-tubulin core^2^, triggering its rapid depolymerisation (**Fig. 1a**, right). Re-establishment of the stabilising GTP-tubulin cap (rescue) protects the unstable GDP-tubulin core and allows growth to resume. Dynamic instability is an intrinsic property of tubulin. In cells, an array of end-binding factors, polymerases, depolymerases and other tubulin-interactors binds the tips of microtubules and modulates their dynamic instability^3^.

**Figure 1.**
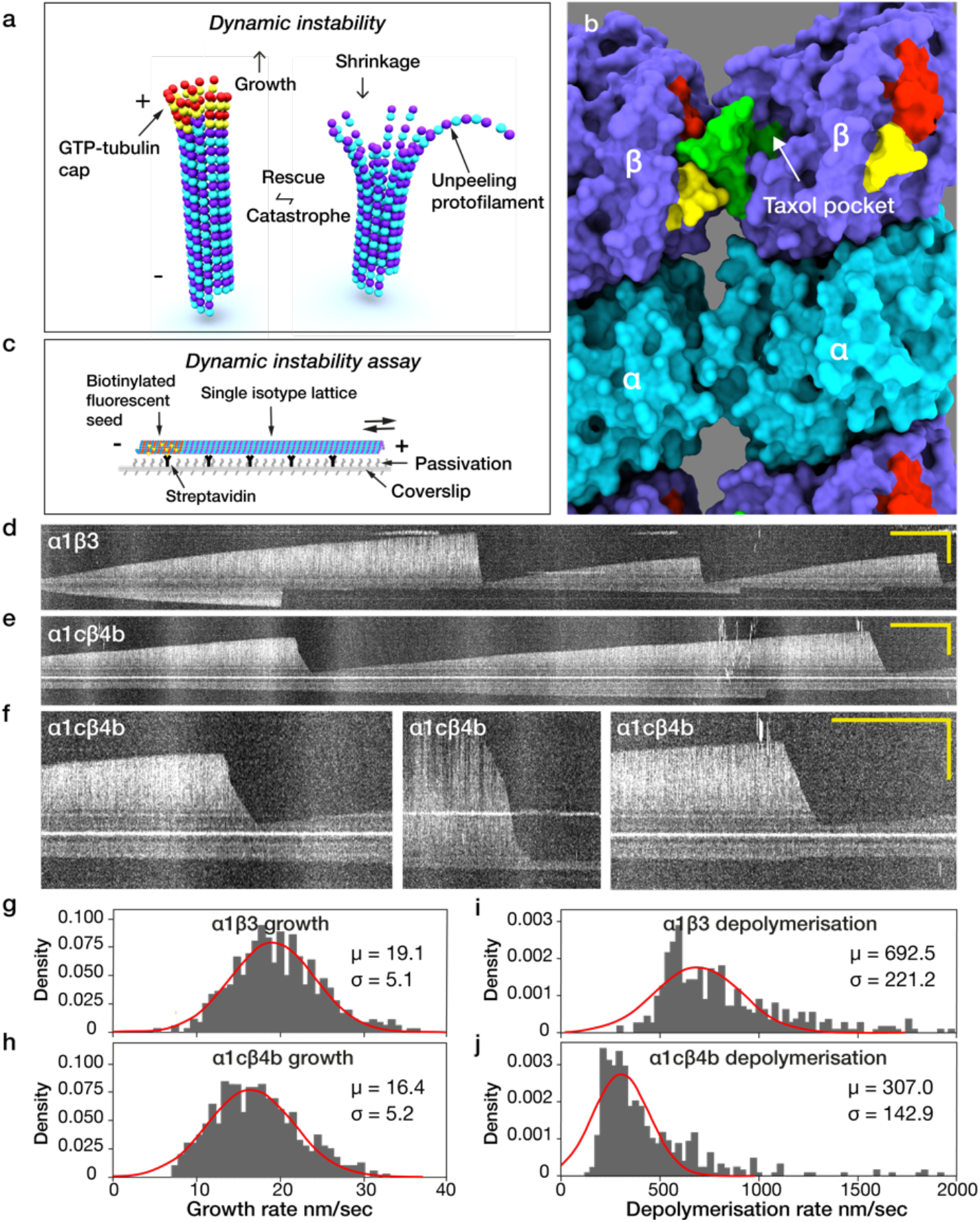
α1β3 and α1cβ4b microtubules depolymerise at different rates. **a**, Dynamic instability. Microtubules grow (left) by addition of GTP-tubulin to their ends, forming a stabilising cap (red & yellow). Loss of this cap (catastrophe) exposes the unstable GDP-tubulin core of the microtubule, which shrinks (right) via rapid loss of curved GDP-tubulin subunits. Re-establishment of the cap (rescue) reverts the microtubule to steady growth. **b**, Molecular graphics showing the taxol binding pocket of β tubulin as viewed from the microtubule lumen. The pocket abuts the M-loop (green, TARGSQQYRA) which links protofilaments in the lattice by engaging the H1-S2 (yellow, AAGNKYVP) and H2-S3 (red, RPDNFVF) loops of the neighbouring subunit. Similar links form between α tubulins (not highlighted). Graphic made in Chimera X^49^, using 6EVW.pdb.**c,** Schematic of flow cell for assays of microtubule dynamics. **d-f,** kymographs showing spontaneous catastrophe and depolymerisation of **d**, α1β3 and **e,f**, α1cβ4b single isotype microtubules. **f**, examples of 2-phase, 3-phase and 1-phase depolymerisation of α1cβ4b lattices. Leftmost and rightmost views are zooms from **d**. Scale bars 10 μm (horizontal) and 2 mins (vertical). **g,h**, comparison of growth rates for **g**, α1β3 and **h**, α1cβ4b microtubules at 12 μM tubulin. Growth rates are similar but statistically different (see Methods). Depolymerisation rates for **i**, α1β3 and **j**, α1cβ4b are clearly different. μ: mean; σ: standard deviation.

Recent work has shown that in addition to classical dynamic instability, dynamic conformational changes in the core GDP-tubulin lattice of polymerised microtubules can also occur, dependent on the binding of allosteric effectors^1,2^. Aside from GTP, GDP.Pi and GDP^3,4^ and their analogues (GMPCPP^5^, GTP<u>γ</u>S^6^), allosteric effectors for tubulin conformation in the lattice include neighbouring tubulin molecules ^7^, a variety of lattice-binding macromolecules, for example EB proteins^8^, doublecortin^8,9^, kinesin molecular motors^10,11^, and a number of lattice-binding drugs^12^. Taxol (generic name Paclitaxel) was the first tubulin-specific drug to be discovered^13^. It is a microtubule-stabilising small molecule effector used in the chemotherapy of breast, lung and ovarian cancer. Taxol binds β tubulin, on the inside (luminal) surface of microtubules, adjacent to the M-loop. The M-loop connects laterally between protofilaments (**Fig. 1b**). Taxol shares its binding site with other taxanes such as docetaxel (Taxotere^®^) and cabazitaxel (Jevtana^®^) and with non-taxanes such as epothilone. Taxol is proposed to directly stabilise lateral connections between protofilaments and to allosterically stabilise the longitudinal interface between neighbouring heterodimers in a protofilament^4,14–16^. Precise molecular mechanisms remain unclear. Exactly how taxol works in cells is also not fully clear. Taxol was initially thought to arrest mitosis by activating the spindle assembly checkpoint, ultimately resulting in cell death. More recently, taxol has been shown to cause multipolar divisions, and thereby aneuploidy and the consequent death of daughter cells with incomplete sets of chromosomes^17^.

In mixed-isotype brain microtubules, taxol not only stabilises the core GDP-tubulin lattice, but also expands the axial spacing between tubulin heterodimers compared to that of the unliganded GDP-tubulin lattice^18^. There are indications that the extent of expansion can depend on whether taxol binds before or after the lattice assembles^19^. Ordinarily, microtubule assembly requires GTP, but taxol can also drive the assembly of GDP-tubulin into microtubules^20^, and can stabilise short protofilaments in solution^21^. Fluorescent analogues of taxol have been shown to create and/or bind preferentially to lattice defects. Defects have their own, local dynamics, allowing new tubulin to be incorporated in a repair process^22^.

Humans have 9 α and 10 β tubulin isotypes^23,24^. It is unclear why humans need so many tubulins, but we do: single point mutations in various human tubulin isotypes are documented to cause specific human diseases (tubulinopathies), often linked to developmental abnormalities^25,26^. Taxol resistance in patients can be accompanied by an increase in β3 tubulin expression^27^, suggesting that β3 tubulin expression might protect against taxol. The possibility that taxol affects different human tubulin isotypes differently has not been directly explored. Here we test this possibility, using microtubules assembled from purified single isotype tubulins. We show that human α1bβ3, zebrafish α1cβ4b and human α1bβ4b microtubules have different lattice stabilities and different responses to taxol. Further, by studying mosaic-isotype microtubules and segmented-isotype microtubules, we probe the spatial range of taxol-dependent conformational switching in the different lattices, with relevance to the molecular mechanism of taxol-induced microtubule stabilisation, and to cancer medicine.

## Results

We chose to examine α1β3 and α1β4b tubulin isotypes. β3 tubulin is confined largely to neurons^28^, whilst β4b tubulin has a broader tissue specificity^29^. Tissue specificity of α tubulin isotypes is less well-understood. The primary sequences of human α1bβ3 and zebrafish α1cβ4b tubulins differ at 39 positions (28 outside of the C-terminal E-hooks). Zebrafish α1cβ4b tubulin dimer differs from human α1bβ4b by 2 residues in α and 5 residues in β tubulins. Hereafter, “α1cβ4b” implies zebrafish tubulin and “α1bβ4b” implies human one, whilst “α1β4” implies both. To find out if the residue substitutions that delineate these isotypes make a difference to the molecular action of taxol, we assembled defined-isotype lattices and compared their *in vitro* properties and the influence of taxol on those properties. We expressed tubulins in insect cells and used dual affinity tag-purification, with an 8x his tag and a FLAG tag on the C-termini of a and β tubulins, respectively (see Methods and^30^), to ensure we obtained only defined-isotype full length tubulin heterodimers. Experiments quantifying dynamic instability used dark field illumination of unlabelled tubulins. For kinesin-driven microtubule gliding assays, 5% HiLyte 488- or HiLyte 647-porcine brain tubulin was included as a trace fluorescent label (see Methods). Single isotype tubulins were not fluorescently labelled.

### α1β3 GDP-tubulin lattices depolymerise faster than α1β4 GDP-tubulin lattices

α1β3 and α1β4 single isotype microtubules nucleated from GMPCPP porcine brain microtubule seeds have markedly different dynamics. α1β3 microtubules catastrophise more frequently and depolymerise much faster than α1cβ4b microtubules (**Fig. 1c-f**). In both α1β3 and α1cβ4b microtubules, depolymerisation following catastrophe is usually monophasic, but α1cβ4b microtubules can depolymerise in multiple phases (**Fig. 1e**). Since these are single isotype microtubules, depolymerisation in two or more sequential phases implies a structural difference between the corresponding regions of the lattice, for example a change in protofilament number, occurring at a lattice defect^31^. Growth rates of α1β3 and α1cβ4b microtubules are similar (**Fig. 1g, h**), but α1β3 microtubules shrink around 2-fold faster than α1cβ4b microtubules (**Fig. 1i, j**). Note that we fitted both growth and depolymerisation histograms with gaussians, but for depolymerisation the fits are poor, consistent with multiple types of depolymerisation event (including, for α1β3 lattices, multiple types of monophasic event).

### Taxol differentially stabilises α1β4 versus α1β3 lattices

We find, as has been previously suggested^27^ that α1β3 lattices are relatively insensitive to taxol. 50 nM taxol has no detectable effect on dynamically unstable α1β3 microtubules (**Fig. 2a**). By contrast, 50 nM taxol is highly effective in stabilising α1cβ4b microtubules against depolymerisation, abrogating catastrophe and driving continuous (processive) growth (**Fig. 2b**). Taxol thus can amplify the intrinsic stability difference between α1cβ4b and α1β3 microtubule lattices. For α1β3 microtubules, 250 nM taxol partially inhibits catastrophe, slows depolymerisation and promotes rescue (**Fig. 2c**, blue arrows). 500 nM taxol fully inhibits α1β3 catastrophe and significantly reduces the rate of plus end growth from 19.4 ± 5.3 nm/sec (median ± SD, n = 20) at 12 μM tubulin in the absence of taxol to 15.6 ± 7.2 nm/sec (n=20) in its presence (**Fig. 2d, e**). For minus ends in the absence of taxol (**Fig. 2d**), a population is present for which growth is too slight to measure and is assigned as zero. This population disappears on adding taxol (**Fig. 2e**), reflecting that 500 nM taxol converts both ends of α1β3 microtubules to processive growth. Growth rates of α1cβ4b microtubules were not obtained because the lattice develops kinks at this taxol concentration (see below).

**Figure 2.**
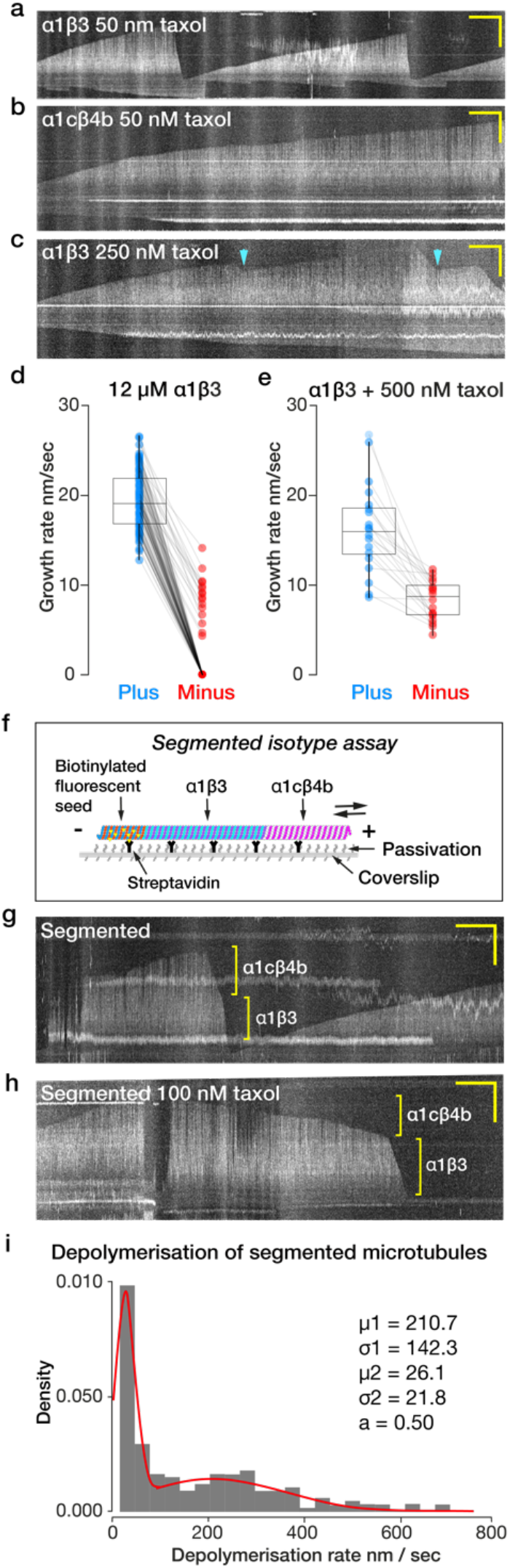
Dynamic instability of α1β3 and α1cβ4b microtubules responds differently to taxol. Kymographs showing **a**, dynamic instability of an α1β3 microtubule in 50 nM taxol, **b**, dynamic instability of an α1cβ4b microtubule in 50 nM taxol. **c**, dynamic instability of an α1β3 microtubule in 250 nM taxol. Catastrophes still occur but depolymerisation is slower, and rescues (cyan arrows) are seen. **d**, **e,** plus end (blue) and minus end (red) growth rates of α1β3 microtubules at 12 μM tubulin **d**, in the absence of taxol and **e**, in the presence of 500 nM taxol. Each dot represents average growth or depolymerisation of individual microtubule. **f**, Schematic of segmented isotype microtubule dynamics assay. **g,h** Biphasic depolymerisation of segmented isotype microtubules **g**, in the absence of taxol. **h**, in 100 nM taxol. Scale bars for all kymographs 2 min (horizontal) and 10 μm (vertical). **i**, depolymerisation rates of segmented isotype microtubules in 100 nM taxol.

### Taxol differentially stabilises α1β4 versus α1β3 segments in segmented-isotype microtubules

To probe for axial communication between adjacent single-isotype regions of the same microtubule lattice, we assembled microtubules made of two contiguous single-isotype segments, an initial α1β3 tubulin segment and a distal α1cβ4b segment (**Fig. 2f**). At 100 nM taxol, α1β3 lattices are susceptible to catastrophe, but rates are somewhat reduced, allowing us to obtain enough initial α1β3 segments of sufficient length. α1cβ4b tubulin was then added and the second segment assembled, whilst maintaining taxol at 100 nM. On washout of free tubulin, whilst maintaining 100 nM taxol, these single-isotype segments depolymerised sequentially in clearly distinct phases, with α1β3 microtubules again depolymerising faster than α1cβ4b (**Fig. 2g**). The transition between phases occurred abruptly, with no detectable tendency of the proximal, intrinsically faster-depolymerising α1β3 segment to accelerate the depolymerisation of the distal α1cβ4b segment close to their intersection. Taxol differentially inhibited depolymerisation of the two segments, broadly in line with its effect on the corresponding single isotype microtubules (**Fig. 2h**). In 100 nM taxol, there is an ~8-fold difference in the depolymerisation rate of α1cβ4b versus α1β3 segments (**Fig. 2i**).

### 500 nM taxol puts kinks into α1β4 microtubules; taxol washout relaxes the kinks

When assembled in 500 nM taxol, α1cβ4b microtubules grow processively and acquire kinks, whereas α1β3 microtubules do not (**Fig. 3a, b**). Even when assembled at 5 μM taxol, kinks were not observed for α1β3 microtubules. At 100 nM taxol, kinks were absent from α1cβ4b microtubules (not shown), suggesting that taxol concentration must exceed a threshold to cause kinking of microtubules. Kinky α1cβ4b microtubules assembled in 500 nM taxol straighten upon washing out taxol using a solution of free tubulin only, showing that kinking is reversible (**Fig. 3c**). Washout with 100 nM taxol in the absence of free tubulin also relaxes kinks. We hypothesise that kinks are produced by localised, taxol-induced expansion of the α1cβ4b lattice, which reverses upon taxol washout (**Fig. 3d**).

**Figure 3.**
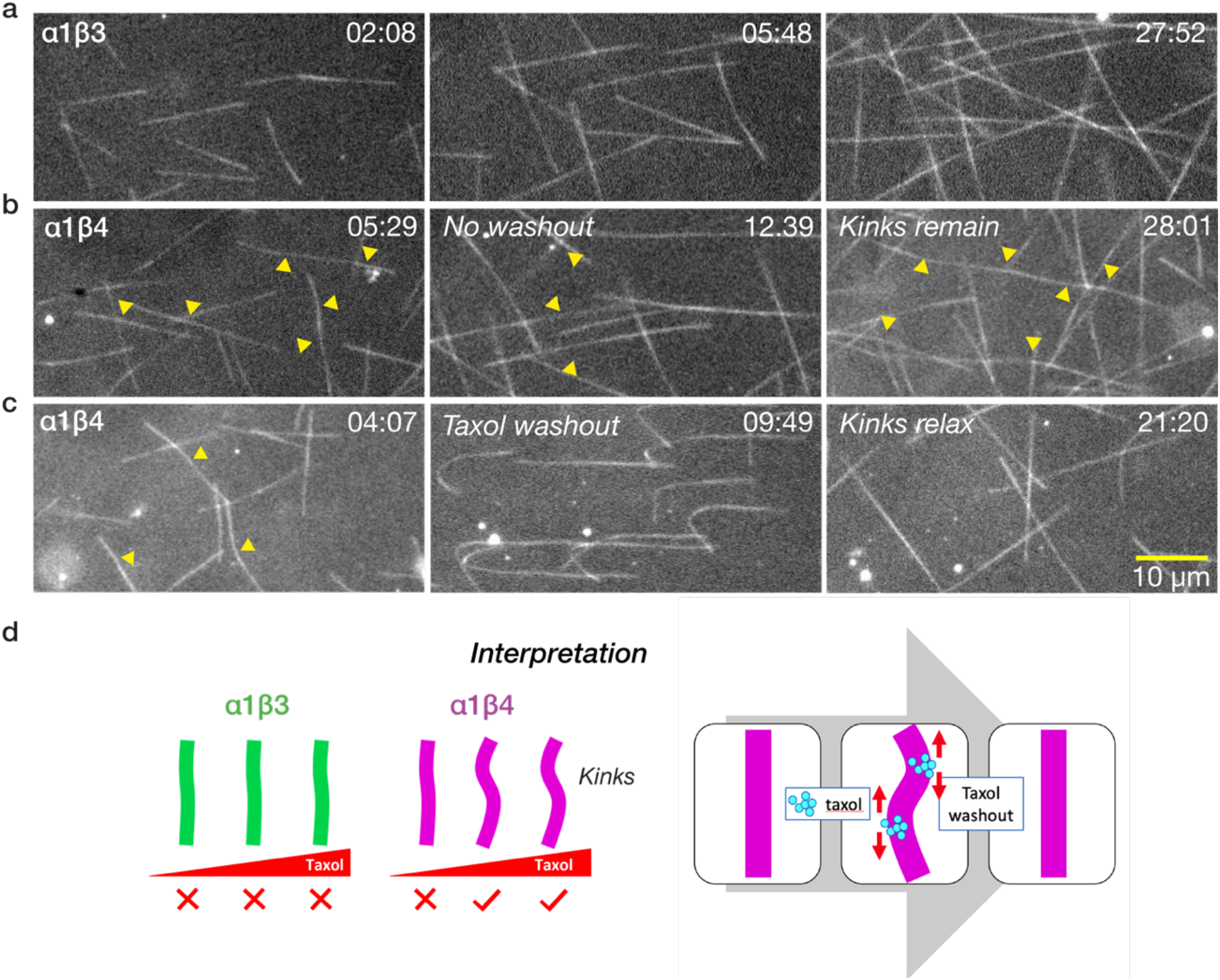
Taxol puts kinks into α1cβ4b but not α1β3 microtubules. **a**, α1β3 microtubules grow processively and straight in the presence of 500 nM taxol at 12 μM tubulin. **b**, under the same conditions, α1cβ4b microtubules develop kinks **c**. Washing out taxol with free tubulin relaxes the kinks. Scale bar 10 μm. Times in min:sec.

### Taxol accelerates the gliding motility of α1β4 but not α1β3 microtubules

Kinesin can sense a taxol-dependent difference in the conformation of α1β3 versus α1β4 GDP microtubules, which emerges as different microtubule gliding rates. At 200 nM taxol, both human and zebrafish α1β4 GDP-microtubules glide over a surface of full-length kinesin-1 dimers at ~450 nm/sec (**Fig. 4a**). Quantification of gliding at 200 nM taxol was less precise because microtubules depolymerise as they glide. Gliding velocity is perhaps 5%-10% under-estimated due to this effect. In 10 μM taxol, both human and zebrafish α1β4 GDP-microtubules accelerate to almost 800 nm/sec, whereas α1β3 microtubules do not (**Fig. 4a**). We could not measure the gliding velocity of α1β3 microtubules at submicromolar taxol concentrations as the microtubules depolymerised too rapidly. For α1β4 microtubules, individual microtubules at any taxol concentration tended strongly to move at either the fast or the slow speed (**Fig. 4a**). GMPCPP microtubules, both α1β3 and α1cβ4b, move at the faster rate (**Fig. 4b**).

**Figure 4.**
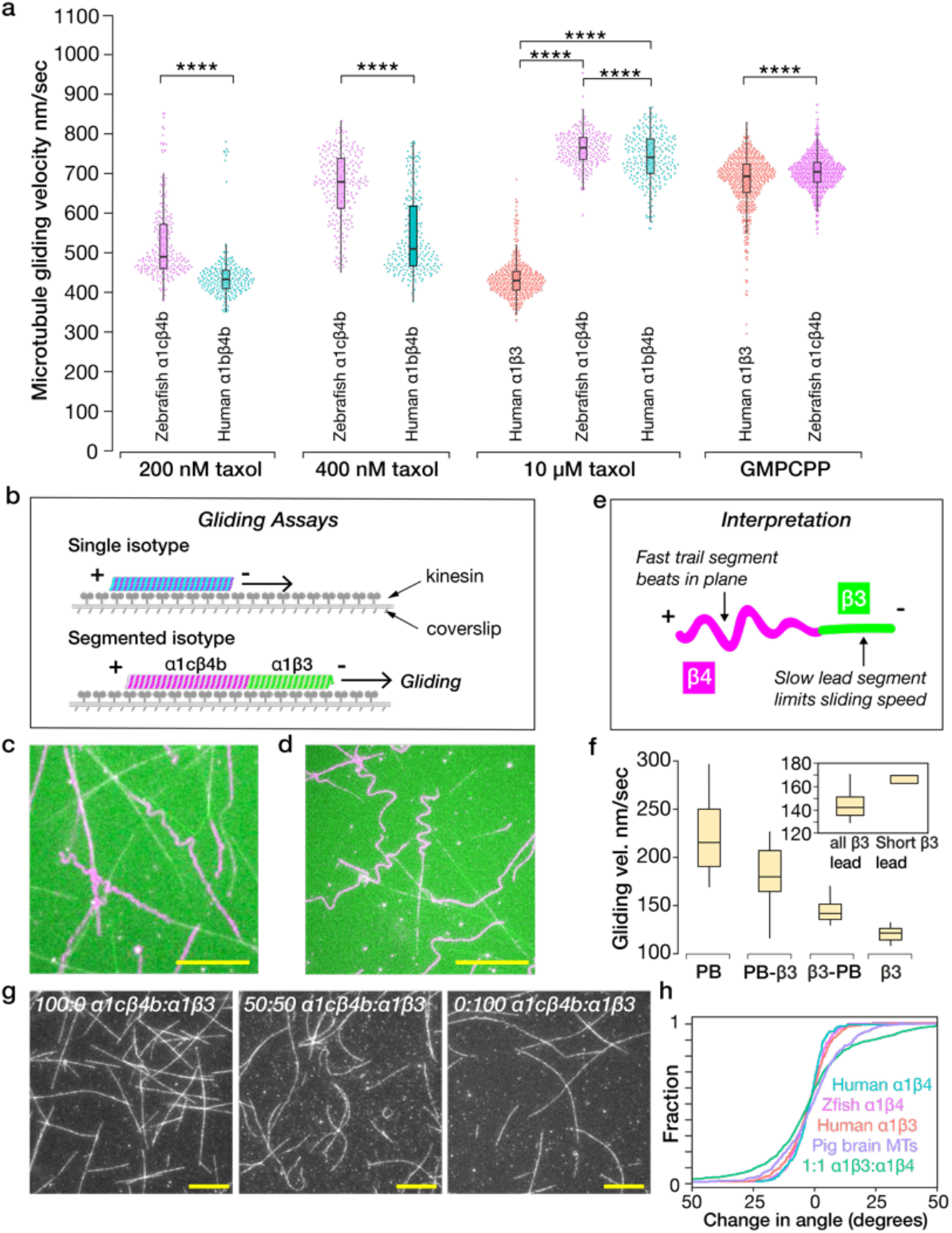
Taxol accelerates kinesin-driven gliding for α1β4 but not α1β3 lattices. **a**, Higher taxl concentrations specifically accelerate α1β4 microtubules, both human and zebrafish. At 200 nM taxol, zebrafish α1cβ4b microtubules glide at 491 ± 96 (n = 231) nm/sec; human α1bβ4b microtubules glide at 434 ± 65 (n = 238). At 400 nM taxol, zebrafish α1cβ4b microtubules glide at 680 ± 87 (n= 219) nm/sec; human α1bβ4b microtubules glide at 511 ± 99 (n = 229). At 10 μM taxol, human α1β3 microtubules glide at 431 ± 49 (n= 505), zebrafish α1cβ4b microtubules glide at 766 ± 42 (n= 242) nm/sec; human α1bβ4b microtubules glide at 742 ± 63 (n = 236), n = instantaneous velocity, see Methods. Quoted velocities are median ± SD. GMPCPP α1β3 and α1cβ4b microtubules both slide at the faster rate at 694 ± 69 (n = 653) and 705 ± 44 (n = 690) nm/sec, respectively. **b,** Schematic of gliding assays. **c,d**, Segmented isotype gliding assays. Trailing α1β4 segments of gliding segmented-isotype microtubules meander behind their leading α1β3 segments. Scale bar is 20 μm. **e,** Schematic. **f**, Gliding velocities of segmented porcine brain-α1β3 isotype microtubules, in 2 mM ATP plus 2 mM ADP: porcine brain segment leading 180 ± 32 (n = 18), porcine brain segment only 216 ± 42 (n = 20), α1β3 segment 142 ± 15 (n = 15) and α1β3 segment only 122 ± 8 (n = 16) nm/sec, velocities are shown as median ± SD, n = average velocity of one microtubule. Inset compares velocities of microtubules with a shorter leading α1β3 segment to velocities of all microtubules with a leading α1β3 segment. **g**, gliding of mosaic isotype microtubules. **h**, Relative change in gliding direction for single isotype versus mosaic isotype microtubules (see *Methods*). A 50:50 mix of α1β3:α1cβ4b appears more sinuous than either single isotype.

### Gliding segmented microtubules ‘beat’ in plane when the α1β3 segment leads

Given that α1β3 and α1cβ4b GDP-taxol microtubules slide at different speeds on kinesin surfaces, we asked what happens to segmented-isotype microtubules gliding on kinesin-1 surfaces (see *Methods*). We find that with the slower α1β3 segment leading (**Fig. 4c**), the intrinsically faster α1cβ4b segment tends to deviate into loops (**Fig. 4d**). Addition of ADP along with ATP makes the loops formed by the α1cβ4b segments larger (**Fig. 4e**), likely reflecting disengagement of some of the kinesins. With a porcine brain tubulin segment trailing behind an α1β3 segment, similar behaviour is seen (not shown). We interpret looping as being driven by the need to balance the net forwards force from the intrinsically faster trailing segment with the net resistive force due to the α1β3 segment (**Fig. 4f**). With an α1β3 segment leading, segmented-isotype microtubules slide slightly faster than do single isotype α1β3 microtubules, suggesting the leading segments are pushed along (**Fig. 4g**), the more so if the leading α1β3 segment is short (**Fig. 4g,** inset).

### Mosaic isotype microtubules glide more sinuously than single isotype microtubules

Gliding mosaic microtubules built from a 50:50 mix of α1β3:α1cβ4b isotypes appear more sinuous than single isotype microtubules of either type (**Fig. 4h,i**) gliding over a kinesin surface in the presence of 10 μM taxol. This might indicate a reduction in bending stiffness, or it might reflect a different pattern of kinesin-generated force.

### Depolymerisation rates of mosaic isotype microtubules depend nonlinearly on isotype ratio

Mosaic microtubules obtained by copolymerising α1β3 and α1cβ4b tubulins blend the properties of the two isotypes, consistent with previous observations on mixing α1bβ1+α1bβ4 and α1aβ3 human tubulin isoforms^32^. Strikingly, depolymerisation rates depend nonlinearly on the α1β3:α1cβ4b isotype ratio in the assembly mix (**Fig. 5a**). Growth rates were similar but significantly different for 100%α1β3 and 100%α1cβ4b tubulins (19.4 ± 5.3 nm/sec and 16.8 ± 5.0 nm/sec, respectively, at 12 μM tubulin, median ± SD) (**Fig. 5b**; same data as **Fig. 1f**).

**Figure 5.**
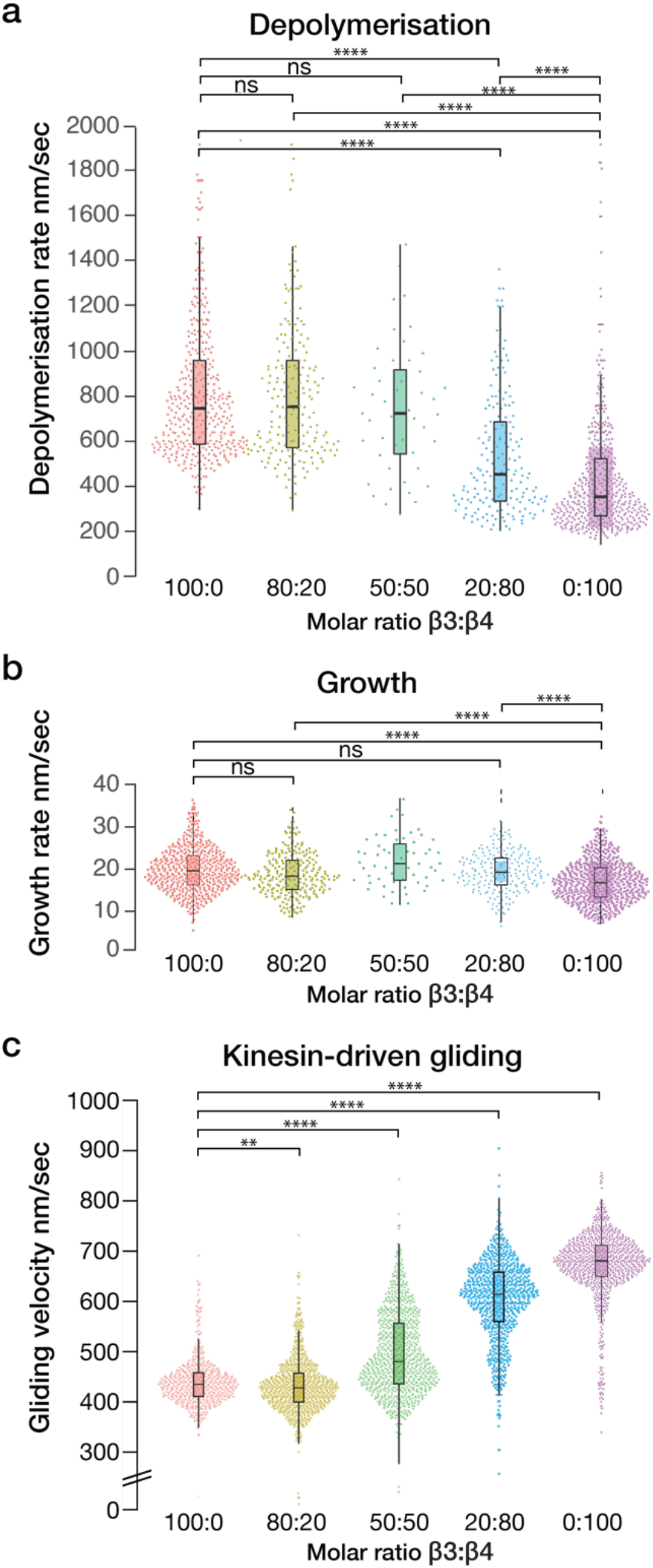
Dynamics and motility of mosaic isotype microtubules depend nonlinearly on isotype ratio. **a**, Depolymerisation rate speeds up markedly with only 20%α1β3 tubulin on a background of 80%α1cβ4b. Depolymerisation rate of microtubules with 100:0 α1β3:α1cβ4b 746.7 ± 324.6 9 (n = 404), 80:20 756.4 ± 357.8 (n = 193), 50:50 724.7 ± 287.2 (n = 47), 20:80 453.3 ± 322.0 (n = 186), 0:100 353.7 ± 277.5 (n = 467) nm/sec (median ± SD). **b**, Growth rates for varying isotype ratios. All experiments performed at 12 μM tubulin, except for the 50:50 ratio, which was at 15 μM tubulin. Growth rate of mosaic microtubules with 100:0 β3:β4 19.4 ± 5.3 (n = 493), 80:20 18.3 ± 5.1 (n = 234), 50:50 21.3 ± 5.8 (59), 20:80 19.3 ± 4.7 (210), 0:100; 16.8 ± 5.0 (n = 551) nm/sec. **c**, Kinesin-driven gliding in 10 μM taxol abruptly switches velocity at a 20:80 α1β3:α1β4 isotype ratio. Gliding velocity of mosaic microtubules with 100:0 β3:β4 431 ± 49 (n = 505), 80:20 424 ± 55 (n = 854), 50:50 476 ± 82 (n = 845), 20:80 609 ± 81 (n = 796) and 0:100 672 ± 69 (n = 950) nm/sec.

### Gliding velocity of taxol-stabilised mosaic isotype microtubules depends nonlinearly on isotype ratio

In mosaic isotype microtubules at 10 μM taxol, the speed of kinesin-driven microtubule gliding also depends nonlinearly on the α1β3:α1cβ4b isotype ratio in the assembly mix (**Fig. 5c**).

## Discussion

We find that taxol affects α1β3 and α1β4 lattices very differently. Taxol inhibits the depolymerisation of both α1β3 and α1cβ4b lattices, but α1β3 lattices require around 10x more taxol than α1cβ4b to fully suppress catastrophe within our ~30 min observation window. Further, taxol accelerates gliding of α1β4 microtubules on surfaces of processive kinesin-1 dimers, whereas α1β3 lattices remain in a slow-gliding conformation even at high taxol concentrations. What are the molecular origins of these different responses of α1β3 and α1β4 microtubules to taxol, and what are their implications?

The differing responses of α1β3 and α1β4 microtubules to taxol show that a small set of mostly conservative-looking residue substitutions (**Fig. 6a**) shifts both the apparent affinity of microtubules for taxol and their ability to execute the taxol-dependent conformational shift responsible for accelerated microtubule gliding. Away from the C-terminal E-hook sequences, most of these residue substitutions are surface-exposed and clustered in the regions of β tubulin that engage the M-loop of the lateral neighbour in the lattice (**Fig. 6b, c**). Crucially, these residue substitutions do not fully inhibit taxol binding to the α1β3 lattice. Taxol at sufficient concentration does stabilise α1β3 microtubules, confirming that taxol does bind to α1β3 microtubules. The clearly different actions of taxol on α1β3 and α1β4 microtubules contrast with the action of GMPCPP, which both stabilises the GDP-tubulin lattice core and accelerates kinesin-driven microtubule gliding, for both α1β3 and α1cβ4b microtubules (**Fig. 4b**). The equivalent effect of GMPCPP on α1β3 and α1cβ4b microtubules demonstrates that both lattices can be shifted into a faster gliding conformation.

**Figure 6.**
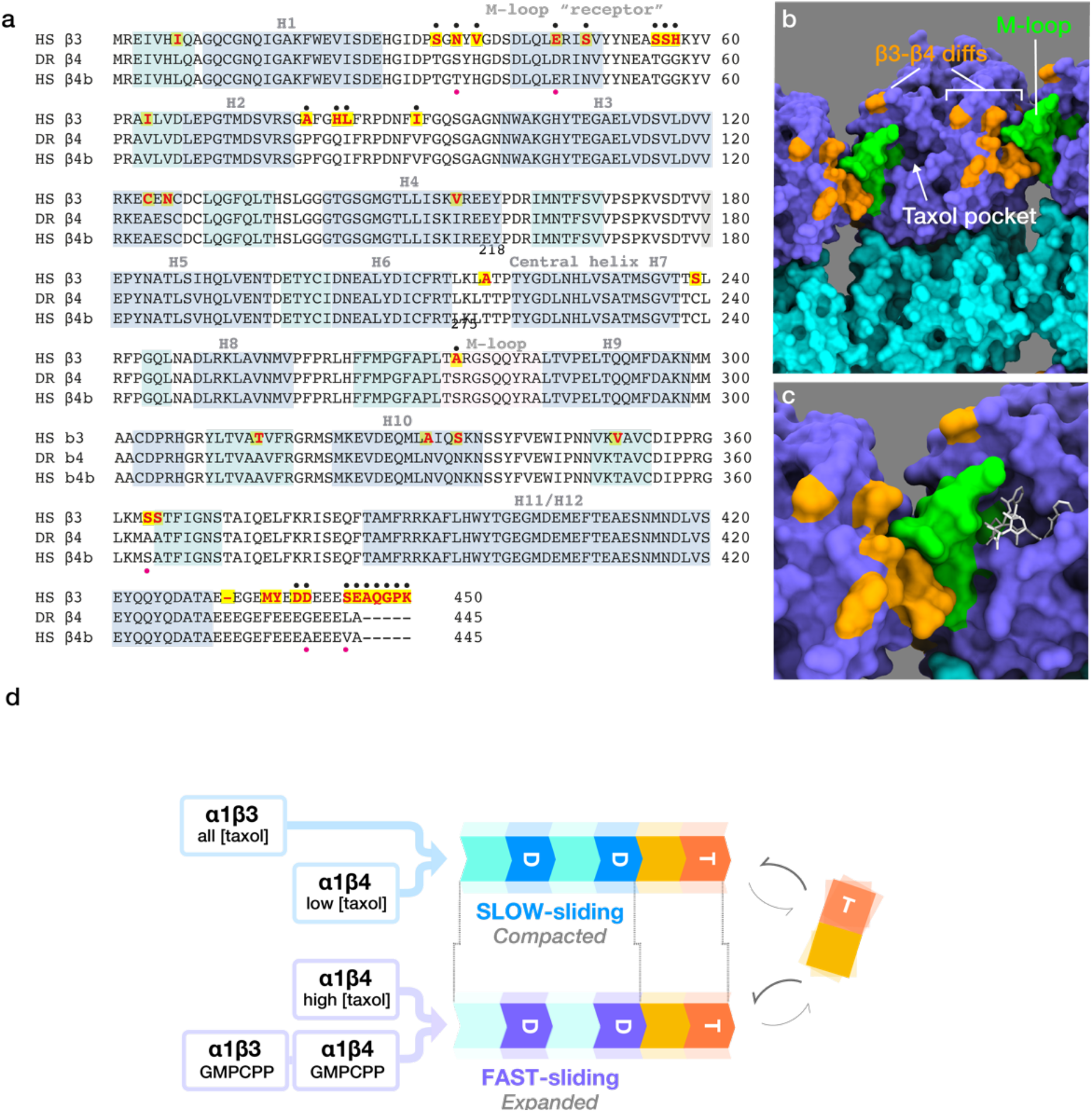
Structural-mechanical differences between β3 and β4 tubulins. **a**, Sequences of human β3, human β4b and zebrafish β4. Helices are shaded magenta, sheets in cyan. Yellow highlighted residues differ between β3 and β4 isotypes. Black dots indicate the subset of these differing residues that is surface-exposed. The M-loop is shaded pink. Red dots highlight the 5 residues that differ between human and zebrafish β4 tubulins. Alignment done using Clustal Omega. **b**, view as in **Fig. 1**, showing the clustering of surface-exposed residue changes between β3 and β4 in the M-loop (green) and in and around the M-loop ‘receptor’ regions (orange). **c**, zoom view of the M-loop region of a GDP-taxol subunit (6WVR.pdb), same colours. **d**, Model. Free GTP tubulin is flexible and has negligibly low affinity for taxol. Incorporation of GTP-tubulin into the growing lattice straightens it as it engages with a lateral neighbour. Hydrolysis and Pi release the GTP-tubulin to GDP-tubulin and introduce strain. Strain in the metastable (strained) GDP-tubulin lattice is relaxed by taxol and other effectors, including GMPCPP. Depending on the effector and the tubulin isotype, effector binding generates either a fast-gliding/ expanded or a slow gliding/compacted conformation of the GDP-tubulin lattice.

Putting these findings together suggests a working model in which taxol binding stabilises both α1β3 and α1β4 lattices but shifts only α1β4 lattices into a faster-gliding conformation (**Fig. 6d**). In this model, the GDP-tubulin lattice can access two effector-stabilised conformational states, with the favoured state dependent on both the effector and the tubulin isotype. The inhibition of α1β3 microtubule plus end growth by taxol (**Fig. 2d, e**) is consistent with previous reports^33^ and suggests taxol reconfigures the α1β3 GTP-tubulin tip-lattice. This point will require further investigation. Our proposed model refers to the GDP-tubulin lattice, and represents a minimal hypothesis in which the GDP-tubulin lattice is shifted between the same two stabilised states by GMPCPP, taxol and other effectors (for example rigor kinesin^10^), even though these stabilising effectors are interacting at widely separated sites on β tubulin. It is possible that other effector-stabilised states of the GDP-tubulin lattice exist; but we only need two to explain our data.

Our model is based on observing microtubule dynamics and microtubule gliding, and not on direct measurement of lattice parameters. Nonetheless, we hypothesise that the transition between slow- and fast-gliding microtubule lattice conformations corresponds to the much-studied transition from a compacted to an expanded lattice. There is robust evidence that both taxol and GMPCPP expand the lattice of mixed-isotype brain microtubules^18^, and that GMPCPP brain microtubules glide faster over kinesin surfaces than GDP brain microtubules^34^. In our own work, we saw that taxol induces kinks in α1cβ4b but not α1β3 microtubules and that taxol washout relaxes the kinks (**Fig. 3**), consistent with them being due to reversible, taxol-dependent local expansion of the α1cβ4b GDP-tubulin lattice. Accordingly, we have named our 2 taxol-stabilised lattice states, SLOW-gliding/Compacted and FAST-gliding/Expanded. The FAST/Expanded state may resemble that of the GTP-tubulin tip-lattice (**Fig. 6**). Our model posits that taxol-dependent stabilisation and taxol-dependent acceleration/expansion are separable. This is evident for both our isotypes. α1β4 lattices are stabilised but not accelerated at low (<400 nM) taxol concentrations, but stabilised and accelerated at higher taxol concentrations. α1β3 is stabilised by taxol but not expanded/accelerated. It is already known that effector-dependent stabilisation and effector-dependent expansion of the microtubule lattice are not necessarily coupled. Prota et al.^35^ recently showed that baccatin III, a taxol precursor, expands but only marginally stabilises brain GDP-microtubules.

How does taxol stabilise both α1β3 and α1β4 lattices, but only accelerate the gliding of α1β4 and not α1β3 microtubules? We see two broad possible explanations for the lack of taxol-acceleration of α1β3. One is that taxol binds and stabilises the core α1β3 GDP-tubulin lattice but that the conformational changes driven by taxol binding do not propagate to the kinesin interface. The other possibility is that the main α1β3 GDP-tubulin lattice binds little or no taxol, but that α1β3 microtubules are stabilised by taxol binding at their ends. Remarkably, there is evidence that at the taxol concentrations used here, only a few percent of taxol sites in the lattice are occupied^36^. This might reflect sparse binding to the lattice, or it might reflect a tendency for taxol to bind preferentially to the microtubule tips, where it can directly influence tubulin subunit exchange, potentially differently for different tubulin isotypes. Taxol binding at microtubule tips might tend to be favoured because the GTP-tubulin tip lattice is expanded relative to the core lattice (**Fig. 6**), and/or because the lumen of the microtubule is more accessible at microtubule tips. Preferential binding of certain fluorescent taxol analogues to tips and to lattice defects has been seen^31^. A taxol analogue, [^3^H]2-m-AzTax, does bind the core β3 GDP-tubulin lattice, but at much-reduced levels compared to other isotypes^37^. Testing whether non-fluorescent taxol can bind the core GDP-tubulin lattice of α1β3 microtubules will require cryoEM imaging of α1β3 at high resolution and high taxol concentrations. If taxol does bind the α1β3 core lattice, our data imply that it does so without causing the go-faster conformational change.

Recent work indicates that taxol binds the GDP-tubulin lattice much more tightly than it binds to free tubulin because the polymerisation switch^38^ reconfigures the M-loop, which otherwise occludes the taxol site^35^. The lateral contacts made by the M-loops may be the dominant determinant of lattice stability^39^. The angles of the lateral bonds made by the M-loop fall as the protofilament number rises, and the resulting strain in the M-loops can be expected to affect their stability^40^. There is evidence that variations in protofilament number can influence the stability of mixed isotype lattices^31^. We see multiphasic, often biphasic, depolymerisation of some α1cβ4b single-isotype microtubules (**Fig. 1**). This must reflect a structural difference between different regions of the depolymerising lattice, possibly differing numbers of protofilaments, flanking a defect. The M-loop sequence is conserved across all human β tubulin isoforms, except for the β3 and β1 (aka βVI) isoforms, both of which have an S275A substitution. β3 tubulin has an additional T218A substitution abutting the M-loop and the central helix H7 (**Fig. 6a**). Computational simulation suggested that T218A restricts the accessibility of the taxol pocket^37^. Crucially, β3 tubulin also has residue substitutions in its ‘M-loop receptor’ regions, which engage the M-loop of the lateral neighbour in the lattice. Aside from changes in and around the M-loop itself, these are our mechanistic prime suspects, because restructuring and repositioning the M-loop can change the accessibility of the taxol site, the ability of the taxol site to signal its occupancy, and the basal (taxol-free) lattice stability, seen as a ~2.2-fold difference in the depolymerisation rates of unliganded α1β3 and α1cβ4b GDP-tubulin lattices. There are further sequence differences between α1β3 and α1β4 tubulins (**Fig. 6a**), in buried positions. Certain buried residue substitutions can profoundly affect allosteric communication in β tubulin^41^.

In segmented-isotype microtubules, taxol increases the difference in lattice stability between α1β3 and α1cβ4b segments. Neighbouring segments of different isotypes do not detectably affect each other’s stability, arguing that the different properties of our α1β3 and α1cβ4b lattices are not propagated axially more than a micron or so. Similarly, in gliding segmented-isotype microtubules, stabilised by taxol, segments retain their characteristically different gliding speeds, again arguing that their conformational properties do not propagate axially over more than about a micron. By contrast, *local* conformational signalling between neighbouring tubulins in mixed isotype lattices is clearly important. A 50:50 α1β3:α1cβ4b mosaic lattice depolymerises at the same rate as a lattice built from α1β3 alone. A 20:80 α1β3:α1cβ4b mosaic has appreciably faster depolymerisation than α1β4 alone, whereas depolymerisation of an 80:20 α1β3:α1cβ4b mosaic is not detectably different to depolymerisation of a pure α1β3 lattice (**Fig. 5a**). The nonlinearity is clear, indicating that tubulin molecules in the lattice influence their near neighbours. For the moment we are uncertain of the range of this conformational signalling, because we cannot read the local isotype distribution in the lattice – we only know the isotype ratio in the assembly mix and biased recruitment of one isotype over the other is a possibility. In mosaic-isotype microtubules at different taxol concentrations, kinesin-driven gliding tends strongly to occur at either a fast or a slow rate (**Fig. 5c**). Only a few microtubules slide at intermediate rates, suggesting individual microtubules may be shifted by taxol between slow-gliding and fast-gliding conformations in a concerted way.

How kinesin senses the conformational difference between α1β3 and α1β4 GDP-taxol lattices remains unclear. Experiments in which human β3 CTTs (C-terminal tails) were substituted for the CTTs of *Saccharomyces cerevisae* tubulin showed no effect of the human β3 CTTs on kinesin-1 velocity^42^, consistent with kinesin-1 stepping rate depending on the structured core^43^ of tubulin. A tendency for kinesin-1 in vitro to enrich to brain microtubules with an expanded lattice has been reported^11^. Our model (**Fig. 6**) posits that the GDP-tubulin lattice can be shifted between FAST/Expanded and SLOW/Compacted states dependent on the isotype, the effector and the effector concentration. Kinesin-1 can expand the GDP-tubulin lattice, and is known to bind preferentially to the GMPCPP-tubulin lattice, which is expanded. Faster binding of kinesin to an expanded lattice might be the basis of faster processive stepping on taxol-stabilised α1β4 versus taxol-stabilised α1β3 microtubules. Does this happen *in vivo?* In neurons, kinesin-1 is a processive, axon-specific transporter of neurotransmitter vesicles. β3 tubulin is largely neuron-specific. Recent work indicates that downregulating human tubulin β3 boosts kinesin motility in neurons (without taxol)^44^. Perhaps relatedly, we see *in vitro* that a small decrease in the proportion of α1β3 tubulin in α1β3: α1β4 mixed isotype microtubules substantially accelerates kinesin transport in the presence of 10 μM taxol (**Fig. 5c**).

Our data provide direct experimental support for earlier suggestions that β3 tubulin in human cells is poorly responsive to taxol and that isotype switching in favour of β3 tubulin allows human tumour cells to resist taxol^27^. α1β3 tubulin lattices *in vitro* are unaffected by taxol below ~100 nM, and weakly affected below ~500 nM taxol. Above these concentrations, taxol does fully stabilise α1β3 lattices. The isotype composition of individual microtubules *in vivo* remains unknown, but we have shown *in vitro* that taxol can change the conformation of one lattice-resident tubulin isotype and not another, and that the outcome for microtubule stability and motility is non-linearly related to the isotype ratio. It will be important to explore the response of different tubulin isotypes to a wider range of microtubule-targeting drugs, including combinations of taxol and other microtubule-directed drugs^45^. Since taxol acts differently on different human tubulin isotypes, it may be that other tubulin drugs also have preferred isotypes and that this might be exploited therapeutically.

## Acknowledgements

We thank Carolyn Moores for insightful comments, Thilani Babuji for technical support and fellow members of the Cross lab and the CMCB for improving this ms. RAC dedicates this work to Linda Amos, who is much missed. Funded by a Wellcome Investigator Award to RAC [220387/Z/20/Z].

## Methods

Tubulin expression and purification is based on^30^ with modifications.

### Constructs

Individual sequence-optimized α and β tubulin genes (GeneArt), with L21 leader sequence AACTCCTAAAAAACCGCCACC at the 5’ to enhance protein expression, were C-terminally joined to 8x his-(HHHHHHHH) and FLAG-(DYKDDDDK) affinity tags respectively with GGSGG inserted as a linker in between. The genes are α tubulin *(Homo sapiens TUBA1B* encodes NP_006073 or *Danio rerio tuba1c* encodes NP_001098596) and β tubulin *(Homo sapiens TUBB3* encodes NP_006077; *TUBB4B* encodes NP_006079 or *Danio rerio tubb4b* encodes NP_942104).

Individual α and β tubulin genes were cloned into separate pLIB plasmids, both of which were subsequently integrated into one pBIG1 plasmid backbone via Gibson Assembly as described ^46^ whereby expression of each tubulin is regulated by a pH promoter. This pBIG-α-β plasmid was propagated in *Escherichia coli* DH5α cells. The resulting positive clone was integrated into baculovirus genome via Tn7 transposition sites in *E. coli* DH10EmBacY to allow subsequent generation of recombinant baculovirus.

A successfully integrated baculovirus clone was selected through appropriate antibiotics and blue-white screening. The recombinant virus DNA bacmid, was isolated using MidiPrep reagents (without the use of spin column). The DNA was precipitated overnight in isopropanol at −20°C, pelleted the next day at 14, 000 x g for 15 minutes, washed in 70% ethanol and pelleted again. The pellet was air-dried and dissolved in water. The isolated bacmid DNA was stored at 4°C. Successful insertion of tubulin genes was verified by PCR using M13 forward and reverse primers.

**Figure S1.**
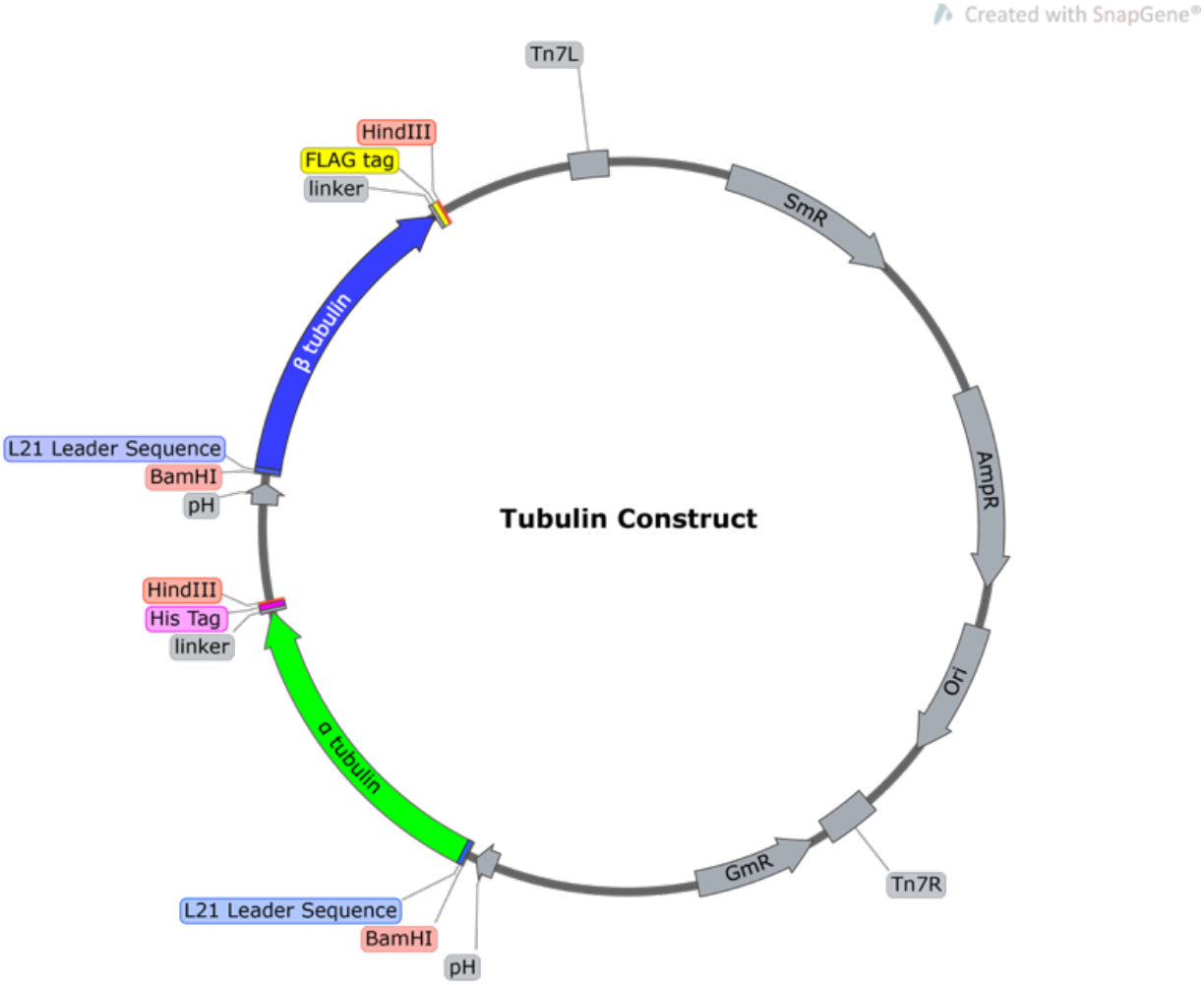
Typical pBIG-α-β plasmid which has Tn7 transposition sites to allow integration of tubulin genes into baculovirus genome.

### Baculovirus generation

Sf9 cells were used for recombinant baculovirus generation and protein expression. Cells were maintained at 28°C with Ex-Cell 420 media (Merck), with shaking at 120 rpm for suspension culture and/or grown as static adherent cells. Generally, cultures were split when reaching a density of approximately 2.0*10^6^ cells/mL to achieve 0.5*10^6^ cells/mL.

For P1 virus production, 2 mL of 0.5*10^6^ cells/mL were seeded onto a 35 mm petri dish and allowed to stand for 15 mins for cell adherence. Transfection mix was prepared by adding 2 μg of bacmid DNA into 200 μL of fresh media, followed by addition of 6 μL of FuGene HD (Promega). This entire mix was dripped onto the culture and incubated without shaking. P1 virus was harvested, typically after 3-5 days of incubation, after clarification by centrifugation at 750 x g for 5 mins at room temperature. P2 virus was generated using one volume of P1 virus added into 100 volumes of fresh culture at a density of 1.0*10^6^ cells/mL. P2 virus was harvested three days afterwards, as for P1 virus, except the supernatant was filtered through a 0.45 μm PVDF syringe filter. P3 virus, which was used for tubulin expression, was generated similarly to P2 virus but 1 volume of P2 virus was used instead to infect 50 volumes of culture. All passages of virus were kept at 4°C for storage, away from light.

### Protein expression and purification

30 mL of P3 virus was added into 1 L of Sf9 culture with a cell density between 1.5 and 2.0*10^6^ cell/mL. Cells were harvested between 54 and 58 hours of incubation at 28°C with shaking at 120 rpm. Cell pellets were collected by centrifugation at 750 x g at 4°C for 20 mins. The pellets were gently resuspended in ice-cold PBS and the cells were collected again by centrifugation. These washed pellets were then flash-frozen in liquid nitrogen and stored at −80°C. On the day of purification, cells were lysed in equal volume of KPEM buffer supplemented with 0.5 M 3-(1-pyridinio)-1-propane sulfonate, ~ 250 unit/mL benzonase, 1 mM DTT, 1 mM PMSF, 0.05% CHAPS, 25 mM imidazole, 1% glycerol, 1 mM ATP and 1 mM GTP. The mixture was homogenised with a douncer with about 60 strokes, then clarified by centrifugation at 200,000 x g (T865 rotor) at 4°C for 1 hr. The supernatant was loaded on to a 5 mL HisTrap HP (Cytiva) at 2 mL/min. After washing the column with KPEM, the protein was step-eluted with KPEM buffer supplemented with 3 mM MgSO_4_, 1 mM GTP and 1 mM ATP, 400 mM imidazole and 250 mM KCl. The pooled fractions were diluted with 2 vols of KPEM buffer to achieve a final concentration of 125 mM KCl and then incubated for 1 hr with 4 mL anti-FLAG monoclonal antibody conjugated resin (M2 agarose, Merck) using rolling. The resulting resin was packed into a 10 mL column, washed with KPEM and eluted with 5 CV of 200 μg/mL FLAG peptide in KPEM supplemented with 125 mM KCl, 1 mM ATP and 1 mM GTP. The pooled eluate was diluted 4 times with KPEM buffer to reduce the KCl concentration to about 30 mM and applied to a 1 mL Capto HiRes Q 5/50 GL column (Cytiva). This column was then washed with KPEM and eluted with KPEM supplemented with 0.5 mM GTP and 0.5 mM ATP, 300-400 mM KCl (isotype dependent). The pooled eluate was exchanged into buffer KPEM without nucleotide using a HiPrep (26/20) desalting column and finally concentrated with an Amicon 30 kD regenerated cellulose spin concentrator to at least 40 μM. Tubulin was snap-frozen in liquid nitrogen in 20 μL aliquots and stored in liquid nitrogen. Aliquots were removed freshly before each experiment.

Concentrations were determined using a spectrophotometer (Cary 50) with molar extinction coefficient 107,110 M^1^cm^-1^ for both *Danio rerio* α1cβ4b and *Homo sapiens* α1bβ4b tubulin and 108 390 M^-1^ cm^-1^ for *Homo sapiens* α1bβ3 tubulin at wavelength 280 nm.

Porcine brain tubulin was purified as described^47^.

*Drosophila* full-length kinesin (gene *Khc;* NP_476590) was C-terminally tagged with 6x his sequence with a thrombin cleavage site in between^48^. Successfully transformed *E. coli* BL21 pLysS cells were maintained at 24°C with shaking at 180 rpm. Due to leaky protein expression, addition of IPTG was omitted which allowed better yield of soluble protein in our hands. Culture was harvested after reaching an optical density of 0.9 at wavelength 600 nm. Kinesin purification was performed at 4°C or on ice unless otherwise stated. The culture was pelleted down at 15,000 x g for 6 minutes. The pellet was gently resuspended in icecold PBS and pelleted down again. Pellets were mixed in 3x pellet mass of buffer containing 10 mM Tris-base, 4 mM Mg-acetate, 250 mM K-acetate, 1 mM TCEP, supplemented with 100 μM ATP, 1x complete protease inhibitor (Roche), 0.5 % vol/vol Triton X-100, ~ 125 unit/mL Benzonase^®^ Nuclease, 0.1 mg/mL lysozyme, with pH adjusted to 8.0 with acetic acid. The bacteria were lysed by sonication (Misonix, S-4000 Ultrasonicator) with 6 cycles of 10-second pulses at 35 Amps, followed by 30-second cooling after every round. This whole cell lysate was clarified at 20, 000 x g for 20 minutes and the supernatant was passed through glass wool. The clarified supernatant was then applied to a 1 mL HiTrap Talon crude column (Cytiva) and eluted with 100-150 mM imidazole in 10 mM Tris-base, 4 mM Mg-acetate, 250 mM K-acetate, supplemented with 1 mM TCEP and 100 μM ATP. The pooled affinity-purified eluates were then further purified by application to a 1 mL HiTrap Q HP column (Cytiva) followed by elution with 200 mM NaCl in 10 mM Tris-base, 4 mM Mg-acetate, 250 mM K-acetate, supplemented with 1 mM TCEP and 100 μM ATP. The pooled eluates were mixed with glycerol to 20% glycerol and snap-frozen in liquid nitrogen for storage.

### Glass slide treatment and flow chamber preparation

Hydrophilic microscopic slides (75*26*1.0 mm, Thermo Scientific) or coverslips (#1.5, Thermo Scientific) were sonicated in 3% Neutracon in ultrapure water (18 MOhms) for 30 mins, followed by 11 rounds of sonications and rinses. The cleaned coverslips were stacked together and stored in a zip-lock bag to minimise contact with air.

### Glass surface passivation for dark-field imaging

Glass slides were plasma-cleaned (Air plasma, Henniker plasma HPT-200) for 5 mins immediately before use. Flow cell channels were assembled using two thin strips of double-sided tape, about 3 mm apart, sandwiched between a glass slide (thickness ~1.0 mm * 75 mm * 26 mm) and a coverslip (no. 1.5, 22 mm * 22 mm). Each channel was preincubated for 30 mins with 0.2 mg/mL of PLL-PEG-biotin (SuSoS), washed out with KPEM, then incubated with 1 mg/mL NeutrAvidin (Thermofisher Scientific) in KPEM buffer, and washed again with KPEM. Biotinylated microtubule seeds were then introduced.

### Microtubule GMPCPP seed preparation

In order to allow one end of microtubule to anchor down to the glass surface, short segments of microtubules called seeds were used. Seed mixture was prepared as 26 μM of porcine brain tubulin containing 10% biotinylated tubulin and 10% HiLyte 488 tubulin (both from Cytoskeleton) in 1 mM GMPCPP in KPEM, incubated on ice for 15 mins, flash frozen in liquid nitrogen and stored over liquid nitrogen. To assemble seeds, an aliquot of seed mixture was thawed and polymerised at 37°C for 30 mins. Residual free tubulin was removed by centrifugation at 20 psi in a Beckman AirFuge for 5 mins at room temperature. Pelleted seeds were gently resuspended by aspiration and diluted in KPEM to obtain optimal decoration of the flow chamber. Unbound seeds were washed out from flow chambers using 1% tween 20 in KPEM.

### Microtubule dynamics assay

Dynamics assays were performed in KPEM buffer, which is 100 mM PIPES, 2 mM EGTA, 1 mM MgSO_4_, pH adjusted to 6.9 with KOH pellets and filtered through a 0.2 μm filter. Tubulin at different concentrations was prepared in KPEM buffer (pH 6.9, 100 mM PIPES, 2 mM EGTA and 1 mM MgSO_4_), supplemented with an oxygen scavenger system (4.5 mg/mL glucose, 0.2 mg/mL glucose oxidase, 35 μg/mL catalase and 0.5% (v/v) β-mercaptoethanol), 1 mM GTP, 1 mg/mL BSA and 0.2% tween 20. All solutions were freshly clarified by AirFuge at 4°C for 5 mins to remove large particles which might interfere with darkfield imaging. To prevent evaporation, flow cell ends were sealed with vacuum grease after tubulin mix was introduced into seed-decorated flow cell channels. To obtain microtubule depolymerisation rates at different taxol concentrations, free tubulin was washed out with taxol solution at a defined concentration. Depolymerisation rates were obtained upon restraightening of microtubules after flow had stopped.

### Darkfield/epifluorescence microscopy

Darkfield Images were acquired with an electron-multiplying charge-coupled device (EMCCD) camera (Andor, iXon DU 897) fitted to Nikon E800 microscope with a 100x objective (Plan Fluor NA 0.5-1.3 variable iris, Nikon). Samples were illuminated with a 100 W mercury short-arc lamp (102D, Ushio) connected to the microscope with a fibre optic light scrambler (Technical video) and the unwanted light was filtered out with a cold mirror and green interference filter with 500-568 nm band pass (Nikon). Epifluorescence was achieved using a stabilised mercury lamp (X-cite exacte, Lumen Dynamics) with light pipe connection to the microscope. Microscope temperature was maintained at 30°C using a custom-made chamber and a heater (Air-Therm ATX, World Precision Instruments). An electronic shutter was used to switch between fluorescence and darkfield illumination. Microscope and camera were controlled by Metamorph software (Molecular Devices). For dynamics experiments, 300 ms exposure and 200 ms exposure time were used for darkfield and epifluorescence respectively, taking 1 frame of epifluorescence for every 300 frames of darkfield illumination, with 1s frame intervals and 160 nm per pixel.

### Microtubule preparation for gliding assay

Motility assays were performed in BRB80 buffer, which is 80 mM PIPES, 1 mM EGTA and 1 mM MgCl_2_, pH adjusted to 6.8 with KOH pellets and filtered through a 0.2 μm filter. Single isoform or porcine brain or mixed isoform tubulin, which was 5% fluorescently labelled with HiLyte 488 or HiLyte 647 conjugated porcine brain tubulin (Cytoskeleton), was assembled at 40 μM at 1 mM GTP for 30 mins at 37°C. Segmented microtubules were prepared by resuspending separately pelleted single isoform microtubules in 10 μM taxol and 1 mM GTP in buffer KPEM and incubating at room temperature overnight to allow end-to-end joining.

GMPCPP microtubules were assembled at 10 μM of tubulin, again with 5% fluorescent tubulin labelling, supplemented with 1 mM GMPCPP in buffer KPEM for an hour at 37°C.

### Microtubule gliding assay

Flow cell channels were assembled from two coverslips (50 mm * 22 mm and 22 mm * 22 mm, thickness no. 1.5) by double-sided tapes (3M Scotch tape). Each channel was about 5 μL. The coverslips were sonication-cleaned as above and coated with either 1% nitrocellulose in amyl acetate by dripping the solution onto a lens tissue and wiping across the surface or coated with 0.2 mg/mL α-casein in BRB 80 buffer supplemented with 1 mM DTT before introducing kinesin into the channel. Kinesin solution supplemented with 1 mM DTT and 0.1 mg/mL α-casein in BRB 80 buffer was introduced into the channel and incubated for 5 mins to allow adsorption to the surface. Subsequently, unbound kinesin was washed out with 5 channel volumes of buffer BRB 80 supplemented with 1 mM DTT and 2 mM ATP (together with or without 2 mM ADP for segmented microtubules). Suitably diluted microtubules, typically about 30-50x dilution in buffer BRB 80 supplemented with 1 mM DTT and 2 mM ATP and 1x GOC with or without corresponding concentration of taxol, was introduced, briefly incubated to allow sufficient microtubule density. Unbound microtubules were washed out with motility buffer with or without taxol accordingly. Fluorescent microtubules were visualised using an eduWOSM microscope. The eduWOSM uses a miniature 4-colour LED light engine and a 100x Nikon oil immersion objective. Please see https://wosmic.org/projects/eduwosm/Index.php for details. Time series were recorded typically at one frame every 5 seconds, taking 200 ms light illumination.

### Data analysis

Kymographs were generated in FIJI using the “MultiKymograph” plug-in, by overlaying an 11-pixel straight-line ROI on microtubules. Growth and depolymerisation rates were determined by fitting a linear ROI over the tip of each microtubule in the kymograph by eye. For multisegmented events, each segment of the ROI was treated as one data point (n = 1), unless otherwise stated. Microtubules with length less than 3 μm were not considered for quantification. For microtubule gliding assays, the Fiji Plugin MTrackJ was used to track the tips of advancing microtubules, taking the angular change due to the most recent displacement vector (the vector pointing from the previous point to the current point of the track). Each instantaneous velocity (determined by every two consecutive frames) was deemed as one data point (n = 1), unless otherwise stated. For statistical analysis we used a Mann Whitney-U test, where ns indicates p ≥ 0.05 and significance is * p < 0.05, ** p < 0.01, *** p < 0.001 and ****p < 0.0001. Fitting of Gaussian distributions was done using R with function non-linear least squares (nls).

